# Duplex PCR assay to determine sex and mating status of *Ixodes scapularis* (Acari: Ixodidae), vector of the Lyme disease pathogen

**DOI:** 10.1101/2024.10.23.619921

**Authors:** Isobel Ronai, Julia C. Frederick, Alec T. Thompson, Prisha Sharma, Michael J. Yabsley, Utpal Pal, Cassandra G. Extavour, Travis C. Glenn

## Abstract

Ticks are a major health threat to humans and other animals, through direct damage, toxicoses, and transmission of pathogens. An estimated half a million people are treated annually in the United States of America for Lyme disease, a disease caused by the bite of a black-legged tick (*Ixodes scapularis* Say) infected with the bacterial pathogen *Borrelia burgdorferi*. This tick species also transmits another six human-disease causing pathogens, for which vaccines are currently unavailable. While *I*. *scapularis* are sexually dimorphic at the adult life stage, the DNA sequence differences between male and female *I*. *scapularis* that could be used as a sex-specific marker have not yet been established. We determine the sex-specific DNA sequences for *I*. *scapularis* (male heterogametic system with XY), using whole-genome resequencing and restriction site-associated DNA sequencing. Then we identify a male-specific marker that we use as the foundation of a molecular sex identification method (duplex PCR) to differentiate the sex of an *I*. *scapularis* tick. In addition, we provide evidence that this molecular sexing method can establish the mating status of adult females that have been mated and inseminated with male-determining sperm. Our molecular tool allows the characterization of mating and sex-specific biology across development for *I*. *scapularis*, a major pathogen vector, which is crucial for a better understanding of their biology and controlling tick populations.

## INTRODUCTION

Ticks have a major health impact globally on humans, livestock, pets and wildlife. A tick vector of high concern for humans in the United States of America is the black-legged tick, *Ixodes scapularis*. This tick species transmits at least seven human-disease causing pathogens (Eisen and Eisen 2018), including the causative agent of Lyme disease (*Borrelia burgdorferi* bacteria), for which nearly half a million people are estimated to be treated annually in the United States (Kugeler et al. 2021).

At the adult life stage, *I*. *scapularis* are morphologically and behaviorally sexually dimorphic. Females are one and half times larger in size than males and have a distinctive reddish brown alloscutum which expands during engorgement (Keirans et al. 1996, Sonenshine and Roe 2014). Notably, adult *I*. *scapularis* females require a full bloodmeal for the completion of oogenesis, whereas males typically do not feed, so are unlikely to transmit pathogens to a host (Kiszewski et al. 2001, Troughton and Levin 2007). In addition, adult females have higher infection rates for *B*. *burgdorferi* compared with males (Hart et al. 2022). Adult female *I*. *scapularis* are therefore a larger public health threat than males.

All ticks have females as the homogametic sex and males as the heterogametic sex, with a variety of sex chromosome cytotypes present across tick species (Oliver Jr 1977, Oliver 1989). *I*. *scapularis* females are XX and males XY, based on cytological studies (Oliver et al. 1993). The Y chromosome is the smallest chromosome of the *I*. *scapularis* karyotype, which includes 13 pairs of autosomes (Chen et al. 1994). Recently, a high-quality scaffolded genome assembly of an individual female *I*. *scapularis* has been generated (De et al. 2023) and the sex pseudochromosomes have been identified (Andrew et al. 2023). However, the sequence differences between male and female ticks that could be used as a sex-specific marker have not yet been established.

Molecular sexing methods have been developed for numerous arthropods. If the heterogametic sex has a hemizygous sex chromosome, the typical strategy for molecular sexing is to detect the presence of a marker (DNA sequence) associated with the unique sex chromosome. This strategy has been used for many arthropod pest species, including the red flour beetle *Tribolium castaneum* (Lagisz et al. 2010), the codling moth *Cydia pomonella* (Fuková et al. 2009) and the African malaria mosquito *Anopheles gambiae* (Krzywinski et al. 2004). A molecular sexing method would be helpful for investigating the biology of *I*. *scapularis* when sex cannot be easily determined morphologically, such as cases of a damaged specimen or the immature life stages.

A secondary use of arthropod molecular sexing methods is determining the mating status of adult females. If a male-specific marker is detected from the DNA of an adult female this suggests she has been inseminated with male-determining sperm and has mated. Using a sexing method to determine mating status has been used for major vector species, including the *A*. *gambiae* (Ng’habi et al. 2007). As up to 70% of adult female *I*. *scapularis* mate off-host (Yuval and Spielman 1990, Kiszewski and Spielman 2002), a high throughput molecular method to assess the mating status of *I*. *scapularis* females would be valuable.

A molecular sexing method that targets male-specific markers has not yet been reported for *I*. *scapularis*. In this study we use resequencing and restriction site-associated DNA sequencing data to identify a male-specific DNA sequence for this species, which is then used for molecular sex identification (duplex PCR).

## METHODS

### Biological samples for resequencing

To identify *I*. *scapularis* sex-specific sequences we first surveyed the genetic diversity of males and females in populations from the four quadrants of this species’ geographic range in the eastern half of the United States (Eisen and Eisen 2023): Rhode Island (Washington County), South Carolina (Aiken County), Louisiana (Rapides County) and Wisconsin (Monroe County). In addition, we included males and females from a unique genetic cluster of *I*. *scapularis* in Florida (Osceola County) that we have previously identified (Frederick et al. 2023). For each of these five populations we collected unengorged, adult ticks (N = 4 of each sex, per population). We confirmed species identity and sex using external morphology (Keirans and Litwak 1989). Samples were stored in 70% ethanol at room temperature.

### Resequencing library preparation, data cleaning and mapping

To obtain whole genomes of individual adult males and females from the five populations, we used a low coverage resequencing approach. DNA was isolated from each tick of the five populations following (Frederick et al. 2023), with the modification that the DNA was size selected using Speed-Beads at a 0.8x Speed-Beads:DNA ratio to remove the small, low-quality DNA fragments. The quality of the size selected DNA was assessed on an agarose gel.

We created resequencing libraries for each sample using a modified half reaction protocol for the NEBNext Ultra II FS DNA Library Prep Kit for Illumina (New England BioLabs Inc.), with a fragmentation time of five minutes, and the unique dual-indexed library approach of Adapterama I (Glenn et al. 2019). The quality of the libraries was checked on an agarose gel. The libraries were cleaned using 1.25x Speed-Beads and DNA was quantified on a Qubit spectrophotometer (Thermo Fisher Scientific). We removed the worst quality (based on gel band brightness and library concentration) male and female library for each population, resulting in 30 libraries (N = 3 of each sex, 5 populations, Table S1).

We pooled the individual tagged libraries based on the DNA concentration and sequenced them on an Illumina NovaSeq (North Carolina State University, Raleigh, NC) S4 PE150 kit, targeting a minimum of 8x raw sequence coverage for all samples. The Genomics Sciences Laboratory at North Carolina State University demultiplexed the pooled samples using iTru5 and iTru7 barcodes we provided for each sample. The sequencing and demultiplexing provided an average of 92,097,751 raw read pairs per individual (min = 61,480,376; max = 183,393,550, Table S1), and an average raw sequencing coverage of 12x (min = 8x; max = 23.9x, Table S1). The raw estimated coverage for each sample was calculated using 2.23 Gb as the genome size (De et al. 2023), and 300 bp as the sequence length due to the PE300 reads.

The sequences were cleaned using Trimmomatic v0.39 (Bolger et al. 2014), with the following parameters: ILLUMINACLIP to remove tags within the TruSeq3-PE-2.fa file; allowing 2 mismatches within tags; an accuracy of 30 for palindrome clipping; an accuracy of 10 for simple clipping, the palindrome alignment having a minimum overlap of 2; and requiring both reads to be kept (ILLUMINACLIP:TruSeq3-PE-2.fa:2:30:10:2:TRUE). We further trimmed the sequences using a sliding window of five bases and a quality of 20 (SLIDINGWINDOW:5:20), and a minimum sequence length requirement of 50 base pairs (MINLEN:50). Cleaning and filtering by Trimmomatic removed an average of 12.5% of read pairs (Table S1), mostly due to small inserts.

Trimmed, cleaned sequences were mapped to the current female *I*. *scapularis* reference genome assembly (GenBank: GCA_016920785.2, ASM1692078v2; (De et al. 2023)) using the very-sensitive parameter in Bowtie2 v2.4.5 (Langmead and Salzberg 2012). Sequences that were unmapped or not the primary alignment, were removed from the bam file using SAMtools v1.10 (Danecek et al. 2021). After mapping, 87.7% of the clean reads were retained, which was 76.7% of raw reads (Table S1). For the filtered bam file, we calculated alignment statistics for each sample using the pileup.sh function from BBMap v38.93 (Bushnell 2014) and then we used BBMap to calculate the coverage per sample after mapping, which was the average across all scaffolds in the genome. Based on the mapped reads, female samples had an average coverage of 7.7x and male samples 6.2x (Table S1).

### Triple-enzyme restriction-site associated sequencing libraries

As the whole-genome resequencing dataset is low coverage and could not be used to identify short or single nucleotide variants between sexes, we supplemented this dataset with triple-enzyme restriction-site associated sequencing (3RAD). We used our previously generated 3RAD dataset of adult *I*. *scapularis* (SRA: PRJNA852262 (Frederick et al. 2023)), which included both the ticks used in the resequencing dataset and ticks collected from additional populations in the United States. If these populations consisted of all males or females, the population was excluded. In addition, we sub-selected so that there were the same number of male and female samples represented per population. This subset (N = 82 of each sex) from 25 populations (Table S2) was used for the subsequent analyses. The sequences were cleaned using process_radtags from Stacks v2.5, providing the restriction sites (EcoRI and Xbal) and internal tags along with parameters (-c) removing any reads with an uncalled base, (-q) trimming low quality bases using the default setting of a sliding window with a raw phred score of 10, and truncating the reads to 140bp (-t) (Catchen et al. 2013).

### Male-specific DNA sequence identification using resequencing and 3RAD datasets

Our bioinformatics workflow to identify male-specific *I*. *scapularis* DNA sequences is detailed in Figure 1.

**Figure 1.**
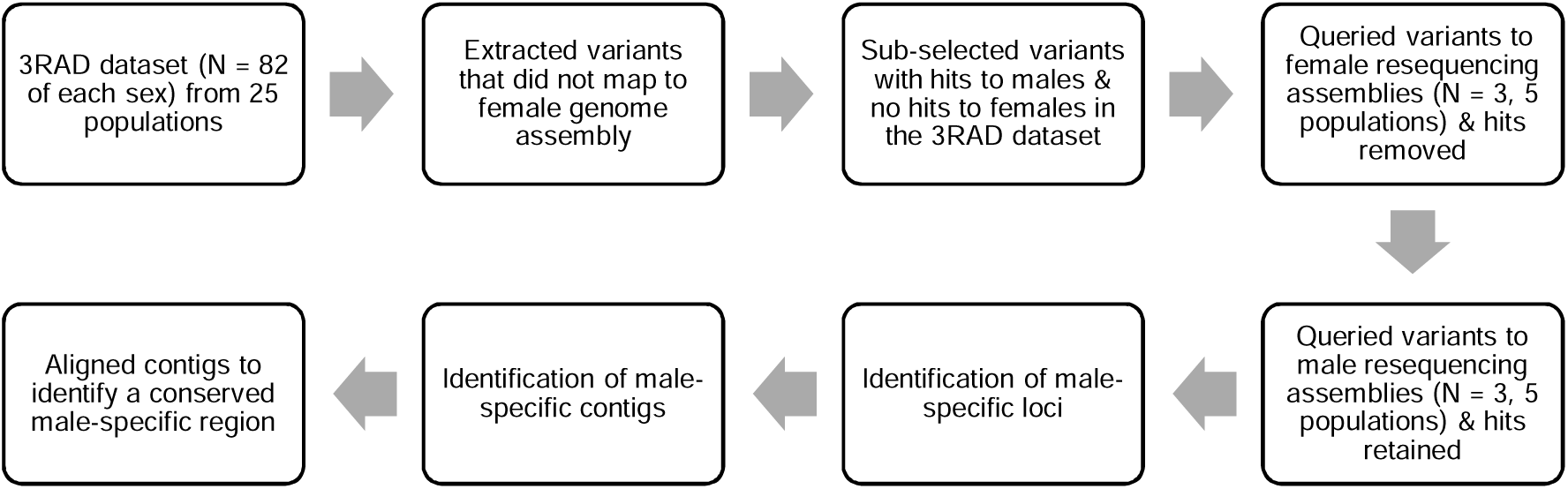
Bioinformatics workflow used to identify male-specific DNA regions of *Ixodes scapularis*.

From the 3RAD dataset we identified small variant sites between males and females. We used the denovo_map.pl program in Stacks, with -M 4 and -n 5, representing the number of mismatches allowed between stacks within an individual and between stacks amongst individuals, respectively. The Stacks populations program was then run using sample sex as the population assignment with parameters -r 0.2, -p 1, then outputting fasta loci, fasta samples, fasta samples raw, vcf, and fstats. We mapped the stacks of sequences from the fasta-samples-raw file to the *I*. *scapularis* genome assembly (GenBank: GCA_016920785.2, ASM1692078v2; (De et al. 2023)) using BWA-MEM v0.7.17 (Li and Durbin 2009). As this genome assembly is from an adult female, we used this assembly as a negative comparison, so stacks that were not mapped to this genome were pulled as potential male-specific loci. These unmapped stacks were then analyzed to determine how many samples were represented by each stack. The potential male-specific loci were sub-selected to stacks that had more than 20 samples represented and consisted of only male samples (i.e. no females). The resultant stacks were checked via Megablast in Geneious Prime 2022.1 (hereafter, Geneious) to the nr/nt database (Kearse et al. 2012). Stacks that had matches to any sequences in the database were removed from consideration. The remaining potential male-specific stacks were then used as queries to the resequencing dataset.

The resequencing dataset was run through SPAdes to assemble contigs for each individual (Bankevich et al. 2012, Prjibelski et al. 2020). The resultant scaffolds.fasta file was exported into Geneious and a custom BLAST database was created per individual. The potential male-specific stacks from the 3RAD dataset were then compared via Megablast to a random subset of the male and female BLAST databases. We identified two overlapping male-specific loci (Table S3) that received no hits to the female databases, but consistent hits to the male databases. These male-specific loci were then run as a BLAST search against each sample’s BLAST database, where they consistently matched male samples with high confidence and either did not match female samples or matched a female with low confidence. We extracted from the sequence files any contigs that matched these male-specific loci. These sequences were then aligned using MAFFT (Katoh and Standley 2013) in Geneious, with the program checking for sequence direction.

We used the male-specific loci to identify a conserved region across the male contigs. For this male-specific region we designed primers (Table 1) using Oligo v6 (Molecular Biology Insights Inc.). Our aim was primers of approximately 25 bp that would produce an amplicon between 100 to 500 bp, a male-specific marker.

**Table 1.**
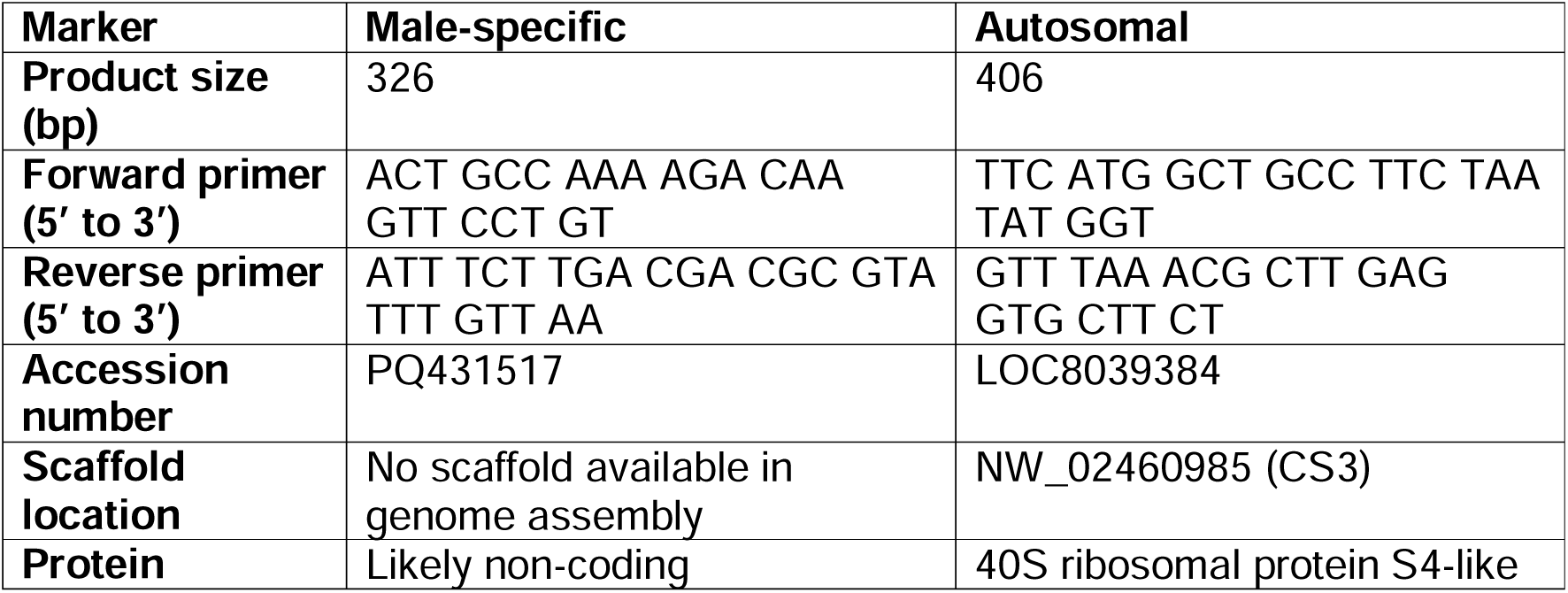
Molecular sexing method (duplex PCR) male-specific marker and autosomal marker primers.

### Cloning of male-specific DNA sequence

The male-specific DNA sequence we identified was inserted into a plasmid to use as a positive control sample for the molecular sexing method. DNA was extracted from a single, lab-reared adult male *I*. *scapularis* that had been morphologically sexed. This tick was deposited by the Centers for Disease Control and Prevention and obtained through BEI Resources, NIAID, NIH: *Ixodes scapularis* Adult (Live), NR-42510. A PCR reaction was conducted following the PCR detailed below, with only the set of primers for the male-specific marker. The amplicon was cleaned using the PCR & DNA Cleanup Kit (Monarch), dA-tailed and cloned using the TOPO TA Cloning Kit for Sequencing (Invitrogen) into the pCR4-TOPO TA vector then One Shot TOP10 Chemically Competent *Escherichia coli* (Invitrogen). The plasmid was purified using the QIAprep Spin Miniprep Kit (Qiagen). The plasmid containing the 326 bp male-specific sequence was deposited with pending (currently available on request from IR).

We confirmed the presence of the amplicon in the plasmid using bidirectional Sanger sequencing (Azenta Life Sciences) and deposited the sequence into GenBank (accession number PQ431517). This sequence was compared to all the *I*. *scapularis* sequences available in the Whole-Genome-Shotgun contigs database using BLASTN (Zhang et al. 2000, Johnson et al. 2008).

### Molecular sexing method (duplex PCR) validation

We designed our molecular sexing method for *I*. *scapularis* as a duplex PCR, a strategy used for other arthropods with heteromorphic sex chromosomes (Fuková et al. 2009, Aguirre et al. 2020). The autosomal marker we used is present in both males and females, the gene *40S ribosomal protein S4-like* (Table 1). We manually designed the autosomal marker primers to be compatible with the optimized PCR conditions for the male-specific marker for the duplex PCR.

We validated the duplex PCR using adults, which were independently morphologically sexed (Figure 2). One adult male and one adult female from a lab-reared colony derived from a northeastern population (Rhode Island), obtained from the Centers for Disease Control and Prevention through BEI Resources. In addition, we tested adults from the genetically distinct southern population of *I*. *scapularis* (Frederick et al. 2023), which have previously been considered as a separate species to the northern population (Oliver et al. 1993), using a lab-reared colony derived from a southern population (Oklahoma), obtained from the Tick Rearing Facility at Oklahoma State University. From each sample we extracted DNA following the method described above, except the DNA was not size selected prior to the duplex PCR. Each DNA sample was diluted to 5 ng/μL.

**Figure 2.**
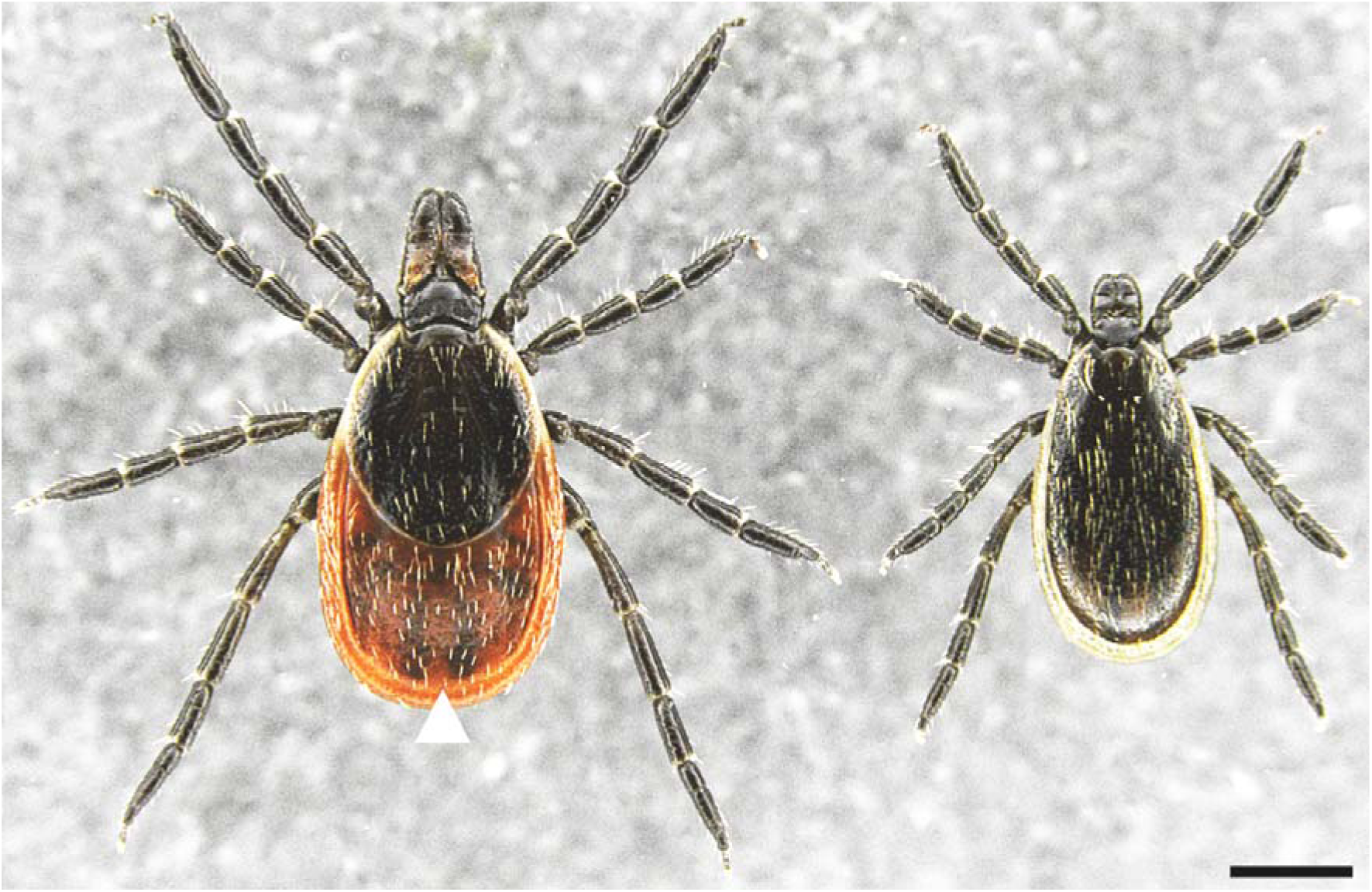
Dorsal view of an adult *Ixodes scapularis* female (left); and male (right). Adult females are larger than males. In addition, adult females have a reddish brown alloscutum that the males lack (arrowhead). Anterior is up and scale bar represents 0.75 mm.

The duplex PCR was conducted in a Biometra TAdvanced High-performance thermal cycler (Analytik Jena) with a reaction volume of 20 μL. PCR reaction: 1X Phusion HFBuffer (New England BioLabs); 200 μM dNTPs; 0.25 μM male marker primers (forward and reverse, Table 1); 0.15 μM autosomal marker primers (forward and reverse, Table 1); 0.4 U Phusion DNA polymerase (New England BioLabs); and 5 ng of template DNA or 0.5 pg plasmid which contains the male-specific sequence (positive control). In addition, the primer set was tested in singleplex at the same concentration. PCR cycling conditions are detailed in Table 2. The PCR products were resolved on a 1.25% agarose gel stained with SYBR Safe DNA Gel Stain (Invitrogen), which included 1 Kb Plus DNA Ladder (Invitrogen).

**Table 2.** Molecular sexing method (duplex PCR) cycling conditions.

| Step | Temperature | Duration | Cycles |
| --- | --- | --- | --- |
| Initial denaturation | 98°C | 30 secs | 1 |
| Denaturation | 98°C | 10 secs | 35 |
| Annealing | 66°C | 30 secs |  |
| Extension | 72°C | 15 secs |  |
| Final extension | 72°C | 5 mins | 1 |

We also validated the sequence of the autosomal marker using bidirectional Sanger sequencing (Azenta Life Sciences), preceded by PCR and gel extraction. We obtained a sequence that aligns to 350bp of the autosomal marker with 91% identity and note that this marker was difficult to sequence, likely due to it being an intronic region that contains nine polyAs.

### Molecular sexing method to determine the mating status of adult females

We investigated whether the molecular sexing method could determine the mating status of adult females. Samples were lab-reared adults obtained from the Centers for Disease Control and Prevention through BEI Resources. Unpaired males (N = 11) and unpaired females (N = 11) were collected in 100% ethanol and stored at −80° C. In addition, we conducted preprandial (off-host) pairings under controlled laboratory conditions, following (Kiszewski and Spielman 2002). The males and females were first frozen (−20° C) for approximately three minutes to increase mating activity. Then single male pairings with one male and one female (N = 12) were placed in a small petri dish (60 x 15 mm) at room temperature (∼22°C). For each pairing we visually observed them as physically attached, with the male being located on the underside of the female (Figure S1) and then the male subsequently detaching from the female. This behavior was initiated within as few as 10 minutes for some pairs, while other pairings took up to two days. Afterwards all paired males were placed in 100% ethanol and stored at −80° C. For the mated females, their eight legs were first removed and the remaining sample (capitulum and idiosoma) placed in 100% ethanol; these were stored at −80° C.

The molecular sexing method was also used to assess the mating status of field collected ticks which had potentially mated. Host-seeking adult females (N = 6) and adult males (N = 6) were collected off-host using drag sampling in Suffolk County, New York. In addition, we collected adult females (N = 3) and adult males (N = 3) from white-tailed deer (*Odocoileus virginianus*) in Delaware (Kent County) and adult females (N = 2) and adult males (N = 2) from American black bears (*Ursus americanus*) in Maryland (Allegany County).

From each tick sample, we extracted DNA following the method described above and each DNA sample, excluding the leg samples, was diluted to 5 ng/μL. With the exception of the 10 host-collected ticks that were tested using a singleplex PCR, the PCR reactions and cycling conditions were as above.

## RESULTS

### Identification of a male-specific *I*. *scapularis* DNA sequence

Our bioinformatics analysis identified two overlapping male-specific loci of 140 bp from the 3RAD dataset (Table S3). These loci had matches to resequencing assemblies from 11 males, with contigs of 1,180 bp to 13,334 bp (Figure S2). In addition, the male-specific loci had a low-quality BLAST hit with a resequencing assembly from one female, a contig of 552 bp (Figure S2). The PCR primers we designed (Table 1, Figure S2) target a highly conserved region across all male contigs that encompasses the two overlapping male-specific loci. In addition, each primer has four SNPs when compared to the potential pseudoautosomal region of the female contig. The *I*. *scapularis* male-specific marker is 326 bp.

### DNA sequence of male-specific marker

We independently validated the male-specific DNA sequence by Sanger sequencing the amplicon of the male-specific marker from a lab-reared adult male. The BLASTN search for the sequence of the amplicon found a match, differing by a single nucleotide polymorphism (position 185: A to C) with unplaced scaffold UN00146945_1, from a whole genome shotgun sequence of *I*. *scapularis*, which is derived from an adult female and pool of adult males (Andrew et al. 2023). In addition, there was an identical match with PKSA02011929.1, from a whole genome shotgun sequence of the embryonic 6 (ISE6) cell line which is derived from pooled embryos (Miller et al. 2018) and a match differing by two single nucleotide polymorphisms (position 185: A to C; and position 234: A to G) with ABJB010723460.1, from a whole genome shotgun sequence of the strain Wikel colony, which is derived from pooled embryos (Gulia-Nuss et al. 2016). There was no match with the current *I*. *scapularis* genome reference assembly (De et al. 2023), which is derived from a single adult female.

### Molecular sexing method (duplex PCR)

The duplex PCR design yields a double band (406 bp and 326 bp) for male samples and a single 406 bp band for female samples (Figure 3). In addition, the positive control plasmid we generated produces a single 326 bp band (Figure 3).

**Figure 3.**
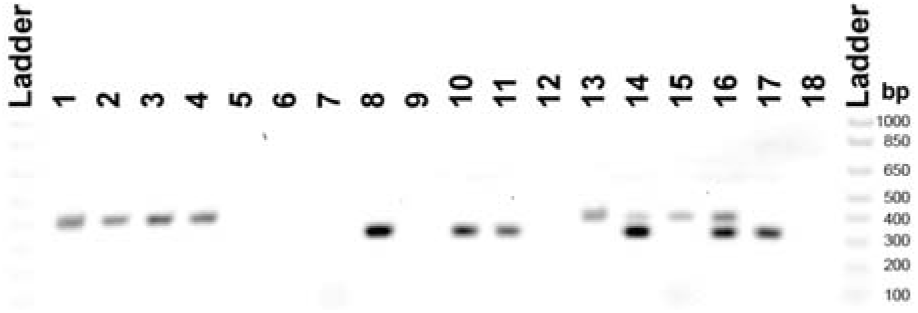
Molecular sexing method (duplex PCR) for *Ixodes scapularis* visualized on an agarose gel; primers described in Table 1. Autosomal marker primer set (Lanes 1-6): female samples have a 406 bp band; male samples have a 406 bp band; and positive control plasmid has no band. Male-specific marker primer set (Lanes 7-12): female samples have no band; male samples have a 326 bp band; and positive control plasmid has a 326 bp band. Two markers in duplex (Lanes 13-18): female samples have a single 406 bp band; male samples have double bands (406 bp and 326 bp); and positive control plasmid has a single 326 bp band. Genomic DNA from *I*. *scapularis* female (Lane 1, 3, 7, 9, 13 and 15) and male (Lane 2, 4, 8, 10, 14 and 16) sourced from the northeastern population at the Centers for Disease Control and Prevention (Lane 1, 2, 7, 8, 13 and 14) or the southern population at the Oklahoma State University (Lane 3, 4, 9, 10, 15 and 16). Positive control plasmid (Lane 5, 11 and 17). No template negative control (Lane 6, 12 and 18).

We successfully determined the sex of 48 male and 48 female adult ticks (Table S4). Our samples include the genetically distinct northern and southern populations of *I*. *scapularis*, indicating that the molecular sexing method is not population-specific.

### Molecular sexing method determines the mating status of adult females

In the laboratory mating assay, we dissected one of the 12 paired females and observed the endospermatophore (Feldman-Muhsam and Borut 1983, Matsuo et al. 1998, Kiszewski et al. 2001) inside this female (Figure S3), which is not present in unpaired females.

We detected the male-specific band in 5 of 11 paired female samples (without legs) from the laboratory mating assay (Figure 4). Whereas, we did not detect the male-specific band in the leg samples from these 11 paired females (Figure 4B). Nor did we detect the male-specific band from any of the whole-body samples of the 11 unpaired females (Figure S4). In addition, we detected the male-specific band in 5 of 6 off-host adult female field samples (Figure S5).

**Figure 4.**
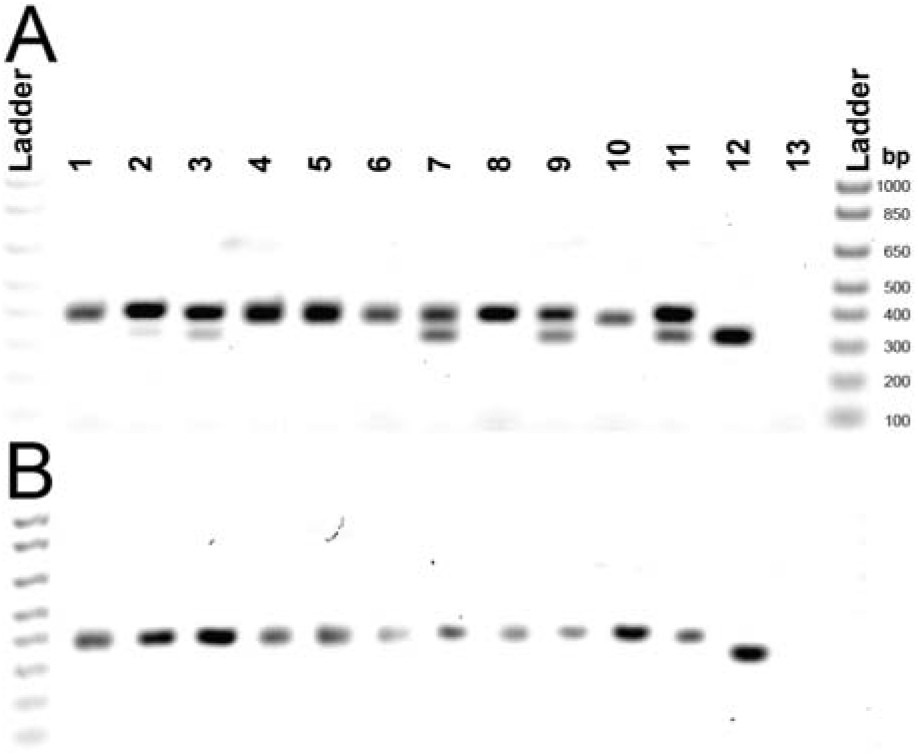
Mating status of adult female *Ixodes scapularis* that were paired with males using the molecular sexing method (duplex PCR) visualized on an agarose gel. Primers described in Table 1. Eleven paired females (Lane 1-11), positive control plasmid (Lane 12) and no template negative control (Lane 13). (A) Paired females (without legs) used as template. All samples have the expected single 406 bp band, however, five samples have a second band at 326 bp (Sample 2, 3, 7, 9 and 11). The positive control plasmid has the expected single band at 326 bp. (B) Legs of the paired females used as template. All samples have the expected single 406 bp band and the positive control plasmid has the expected single band at 326 bp.

## DISCUSSION

We have developed a molecular sexing method (duplex PCR) to identify male and female *I*. *scapularis* using a male-specific DNA sequence. This method is feasible with small tissue samples, we successfully sexed ticks using only the eight legs. Future studies can now investigate how the sex of immature ticks affects biological traits relevant to their role as a pathogen vector, including microbiome, survivability, blood-meal intake, host-seeking behavior and host-preference. Although, *I*. *scapularis* has the most extensive molecular resources for any tick species (Sharma et al. 2022, De et al. 2023), sex-specific DNA sequences that could be used as a sex-specific marker were not yet established. Therefore, the development of a molecular sexing method for this species can be the foundation for insights into the unique biology of ticks. Whether our molecular sexing method can be used for other tick species, given that a male heterogametic XY system occurs throughout the *Ixodes* genus (Oliver Jr 1977, Oliver 1989), remains to be determined.

A secondary use for our duplex PCR is determining the mating status of adult females, as *Ixodes* ticks can mate either on or off-host (Kiszewski et al. 2001). We found that our method is sufficiently sensitive to detect male-determining sperm within paired adult females under controlled laboratory conditions. We also observed male-female pairings that exhibited mating behavior in the laboratory, but for which the duplex PCR did not detect the male-specific marker, suggesting that mating behavior is not a reliable proxy for the insemination status of females. In these pairings, it is likely that the male did not insert a spermatophore and inseminate the female, which has previously been reported in laboratory mating experiments (Kiszewski et al. 2001). We also used the duplex PCR to identify field-collected adult females that likely mated. Future studies will therefore be able to use our duplex PCR to determine the mating status of adult female ticks in circumstances such as surveillance monitoring.

The *I*. *scapularis* male-specific marker is likely located on the Y chromosome. We found the male-specific marker sequence in the recent *I*. *scapularis* genome assembly derived from both sexes (Andrew et al. 2023) and in pooled tick samples that contain both sexes (Gulia-Nuss et al. 2016, Miller et al. 2018), but do not find the marker sequence in the female-only genome assembly (De et al. 2023). The exact genomic location of the male-specific marker remains to be determined, as the *I*. *scapularis* genome assembly that has the sex pseudochromosomes identified, places this marker on an unplaced scaffold (Andrew et al. 2023).

Identifying additional male-specific regions for *I*. *scapularis* may be challenging. Degenerated Y chromosomes have only small amounts of associated DNA sequence and *I*. *scapularis* Y chromosome is less than half the physical size of the X chromosome (Chen et al. 1994). In addition, Y chromosomes are typically gene-poor and composed of more repetitive elements than autosomes (Shaw and White 2022), the *I*. *scapularis* Y chromosome is expected to be mostly repetitive elements as the overall genome is 74% repetitive (Ronai et al. 2024). Lastly, Y chromosomes can be mostly comprised of pseudoautosomal regions with the X chromosome and *I*. *scapularis* Y chromosome is likely mostly pseudoautosomal, as we were only able to identify one male-specific region using our sex-based bioinformatics work flow. Future studies should focus on characterizing the pseudoautosomal regions of the two sex chromosomes in *I*. *scapularis* to help understand the evolution of the sex chromosomes in ticks.

We designed the duplex PCR so that potential PCR issues and contamination can be easily identified. The inclusion of an autosomal marker means that false negatives due to issues with the DNA sample preparation or the PCR reaction are readily identifiable (Aguirre et al. 2020). In addition, if male DNA contaminates a female sample, there will potentially be an anomalous banding pattern, with the male-specific marker (the smaller band at 326 bp) being at a lower concentration than the autosomal marker (the larger band at 406 bp). Therefore, our duplex PCR design provides a robust molecular sexing method.

In conclusion, our identification of sex-specific DNA sequences and development of a molecular sexing method is an important addition to the toolbox for understanding the fundamental biology of *I*. *scapularis* and ticks more broadly. Our study will facilitate future research efforts investigating how sex impacts key traits of a major pathogen vector in the United States, ultimately contributing to better control of tick populations.

## DATA AVAILABILITY

Resequencing data are available under the BioProject: PRJNA853920, SRR19894962-SRR19894991. The plasmid containing the male-specific sequence has been deposited with pending (currently available on request from IR).

## ACKNOWLEDGEMENTS

We thank everyone who provided tick samples, including our research collaborators (Ryan Smith, Jean Tsao, and Risa Pesapane) and community volunteers (Ted Nixon, Tim Lewis, Joe Lloyd, Jeanne Menard, Russ Hartwell, Alex Brown, Jennifer Williston, and Todd Woida). Also thank you to Hein Sprong and colleagues for discussions about *Ixodes ricinus*. We thank Kaylin Chong and The Museum of Comparative Zoology at Harvard University for assisting with imaging the ticks. Lastly, thank you to the members of the Extavour lab for their advice on arthropod molecular methods.

JCF and ATT were partially supported by NSF DGE-1545433 and the IDEAS program at the University of Georgia. JCF was partially supported by NSF IOS-1754950. MJY and ATT were partially funded through Cooperative Agreements AP19VSCEAH00C004 and AP20VSCEAH00C041, Veterinary Services, Animal and Plant Health Inspection Service, USDA. Additional support was provided by SCWDS member state wildlife agencies provided by the Federal Aid to Wildlife Restoration Act (50 Stat. 917) and through federal agency partners, including the United States Geological Survey Ecosystems Mission Area and the United States Fish and Wildlife Service National Wildlife Refuge System. ATT was partially supported by and through the USDA APHIS NBAF Facility Scientist Training Program. The content is solely the responsibility of the authors and does not necessarily represent the official views of NSF, USDA, any other state or federal agency or funding source. IR was supported by an American Australian Association Scholarship and is currently a Howard Hughes Medical Institute Awardee of the Life Sciences Research Foundation. CGE is an Investigator of the Howard Hughes Medical Institute.

## REFERENCES

Aguirre, C., N. Olivares, and P. Hinrichsen. 2020. An efficient duplex pcr method for sex identification of the european grapevine moth *Lobesia botrana* (lepidoptera: Tortricidae) at any developmental stage. J. Econ. Entomol. 113: 2505–2510.

Andrew, B. N., S. L. Johnathan, B. R. Jeremiah, G.-C. Omar, L. Wenlong, S. Arvind, N. P. Michael, B. Saransh, L. S. Mia, M. Molly, H. Isaac Amankona, Z. Xingtan, C. Y. Won, and G.-N. Monika. 2023. The highly improved genome of *Ixodes scapularis* with x and y pseudochromosomes. Life Science Alliance 6: e202302109.

Bankevich, A., S. Nurk, D. Antipov, A. A. Gurevich, M. Dvorkin, A. S. Kulikov, V. M. Lesin, S. I. Nikolenko, S. Pham, and A. D. Prjibelski. 2012. Spades: A new genome assembly algorithm and its applications to single-cell sequencing. J. Comput. Biol. 19: 455–477.

Bolger, A. M., M. Lohse, and B. Usadel. 2014. Trimmomatic: A flexible trimmer for illumina sequence data. Bioinformatics 30: 2114–2120.

Bushnell, B. 2014. Bbmap: A fast, accurate, splice-aware aligner. United States. https://www.osti.gov/servlets/purl/1241166

Catchen, J., P. A. Hohenlohe, S. Bassham, A. Amores, and W. A. Cresko. 2013. Stacks: An analysis tool set for population genomics. Mol. Ecol. 22: 3124–3140.

Chen, C., U. G. Munderloh, and T. J. Kurtti. 1994. Cytogenetic characteristics of cell lines from *Ixodes scapularis* (Acari: Ixodidae). J. Med. Entomol. 31: 425–434.

Danecek, P., J. K. Bonfield, J. Liddle, J. Marshall, V. Ohan, M. O. Pollard, A. Whitwham, T. Keane, S. A. McCarthy, and R. M. Davies. 2021. Twelve years of samtools and bcftools. Gigascience 10: giab008.

De, S., S. B. Kingan, C. Kitsou, D. M. Portik, S. D. Foor, J. C. Frederick, V. S. Rana, N. S. Paulat, D. A. Ray, Y. Wang, T. C. Glenn, and U. Pal. 2023. A high-quality *Ixodes scapularis* genome advances tick science. Nat. Genet. 55: 301–311.

Eisen, L., and R. J. Eisen. 2023. Changes in the geographic distribution of the blacklegged tick, *Ixodes scapularis*, in the United States. Ticks Tick Borne Dis. 14: 102233.

Eisen, R. J., and L. Eisen. 2018. The blacklegged tick, *Ixodes scapularis*: An increasing public health concern. Trends Parasitol. 34: 295–309.

Feldman-Muhsam, B., and S. Borut. 1983. On the spermatophore of ixodid ticks. J. Insect Physiol. 29: 449–457.

Frederick, J. C., A. T. Thompson, P. Sharma, G. Dharmarajan, I. Ronai, R. Pesapane, R. C. Smith, K. D. Sundstrom, J. I. Tsao, H. C. Tuten, M. J. Yabsley, and T. C. Glenn. 2023. Phylogeography of the blacklegged tick (*Ixodes scapularis*) throughout the USA identifies candidate loci for differences in vectorial capacity. Mol. Ecol. 32: 3133–3149.

Fuková, I., L. Neven, N. Bárcenas, N. A. Gund, M. Dalíková, and F. Marec. 2009. Rapid assessment of the sex of codling moth *Cydia pomonella* (linnaeus)(lepidoptera: Tortricidae) eggs and larvae. Journal of Applied Entomology 133: 249–261.

Glenn, T. C., R. A. Nilsen, T. J. Kieran, J. G. Sanders, N. J. Bayona-Vásquez, J. W. Finger, T. W. Pierson, K. E. Bentley, S. L. Hoffberg, and S. Louha. 2019. Adapterama i: Universal stubs and primers for 384 unique dual-indexed or 147,456 combinatorially-indexed illumina libraries (itru & inext). PeerJ 7: e7755.

Gulia-Nuss, M., A. B. Nuss, J. M. Meyer, D. E. Sonenshine, R. M. Roe, R. M. Waterhouse, D. B. Sattelle, J. de la Fuente, J. M. Ribeiro, K. Megy, J. Thimmapuram, J. R. Miller, B. P. Walenz, S. Koren, J. B. Hostetler, M. Thiagarajan, V. S. Joardar, L. I. Hannick, S. Bidwell, M. P. Hammond, S. Young, Q. Zeng, J. L. Abrudan, F. C. Almeida, N. Ayllón, K. Bhide, B. W. Bissinger, E. Bonzon-Kulichenko, S. D. Buckingham, D. R. Caffrey, M. J. Caimano, V. Croset, T. Driscoll, D. Gilbert, J. J. Gillespie, G. I. Giraldo-Calderón, J. M. Grabowski, D. Jiang, S. M. S. Khalil, D. Kim, K. M. Kocan, J. Koči, R. J. Kuhn, T. J. Kurtti, K. Lees, E. G. Lang, R. C. Kennedy, H. Kwon, R. Perera, Y. Qi, J. D. Radolf, J. M. Sakamoto, A. Sánchez-Gracia, M. S. Severo, N. Silverman, L. Šimo, M. Tojo, C. Tornador, J. P. Van Zee, J. Vázquez, F. G. Vieira, M. Villar, A. R. Wespiser, Y. Yang, J. Zhu, P. Arensburger, P. V. Pietrantonio, S. C. Barker, R. Shao, E. M. Zdobnov, F. Hauser, C. J. P. Grimmelikhuijzen, Y. Park, J. Rozas, R. Benton, J. H. F. Pedra, D. R. Nelson, M. F. Unger, J. M. C. Tubio, Z. Tu, H. M. Robertson, M. Shumway, G. Sutton, J. R. Wortman, D. Lawson, S. K. Wikel, V. M. Nene, C. M. Fraser, F. H. Collins, B. Birren, K. E. Nelson, E. Caler, and C. A. Hill. 2016. Genomic insights into the *Ixodes scapularis* tick vector of Lyme disease. Nature Communications 7: 10507.

Hart, C. E., F. A. Middleton, and S. Thangamani. 2022. Infection with *Borrelia burgdorferi* increases the replication and dissemination of coinfecting powassan virus in *Ixodes scapularis* ticks. Viruses 14: 1584.

Johnson, M., I. Zaretskaya, Y. Raytselis, Y. Merezhuk, S. McGinnis, and T. L. Madden. 2008. Ncbi blast: A better web interface. Nucleic Acids Res. 36: W5–W9.

Katoh, K., and D. M. Standley. 2013. Mafft multiple sequence alignment software version 7: Improvements in performance and usability. Mol. Biol. Evol. 30: 772–780.

Kearse, M., R. Moir, A. Wilson, S. Stones-Havas, M. Cheung, S. Sturrock, S. Buxton, A. Cooper, S. Markowitz, and C. Duran. 2012. Geneious basic: An integrated and extendable desktop software platform for the organization and analysis of sequence data. Bioinformatics 28: 1647–1649.

Keirans, J. E., and T. R. Litwak. 1989. Pictorial key to the adults of hard ticks, family Ixodidae (Ixodida: Ixodoidea), east of the mississippi river. J. Med. Entomol. 26: 435–448.

Keirans, J. E., H. J. Hutcheson, L. A. Durden, and J. S. H. Klompen. 1996. *Ixodes* (*Ixodes*) *scapularis* (Acari: Ixodidae): Redescription of all active stages, distribution, hosts, geographical variation, and medical and veterinary importance. J. Med. Entomol. 33: 297–318.

Kiszewski, A. E., and A. Spielman. 2002. Preprandial inhibition of re-mating in *Ixodes* ticks (Acari:Ixodidae). J. Med. Entomol. 39: 847–853.

Kiszewski, A. E., F.-R. Matuschka, and A. Spielman. 2001. Mating strategies and spermiogenesis in ixodid ticks. Annu. Rev. Entomol. 46: 167–182.

Krzywinski, J., D. R. Nusskern, M. K. Kern, and N. J. Besansky. 2004. Isolation and characterization of y chromosome sequences from the african malaria mosquito *Anopheles gambiae*. Genetics 166: 1291–1302.

Kugeler, K., A. Schwartz, M. Delorey, P. Mead, and A. Hinckley. 2021. Estimating the frequency of Lyme disease diagnoses, United States, 2010–2018. Emerging Infectious Disease 27: 616–619.

Lagisz, M., K. E. Wilde, and K. Wolff. 2010. The development of pcr-based markers for molecular sex identification in the model insect species *Tribolium castaneum*. Entomol Exp Appl 134: 50–59.

Langmead, B., and S. L. Salzberg. 2012. Fast gapped-read alignment with bowtie 2. Nature methods 9: 357–359.

Li, H., and R. Durbin. 2009. Fast and accurate short read alignment with burrows–wheeler transform. Bioinformatics 25: 1754–1760.

Matsuo, T., T. Mori, and S. Shiraishi. 1998. Studies on in vitro extrusion and ultrastructure of the spermatophore in *Haemaphysalis longicornis* (Acari: Ixodidae). Acta Zoologica 79: 69–74.

Miller, J. R., S. Koren, K. A. Dilley, D. M. Harkins, T. B. Stockwell, R. S. Shabman, and G. G. Sutton. 2018. A draft genome sequence for the *Ixodes scapularis* cell line, ise6. F1000Research 7: 297.

Ng’habi, K. R., A. Horton, B. G. J. Knols, and G. C. Lanzaro. 2007. A new robust diagnostic polymerase chain reaction for determining the mating status of female *Anopheles gambiae* mosquitoes. The American Journal of Tropical Medicine and Hygiene 77: 485–487.

Oliver, J. H. 1989. Biology and systematics of ticks (Acari: Ixodida). Annu. Rev. Ecol. Syst. 20: 397–430.

Oliver, J. J. H., M. R. Owsley, H. J. Hutcheson, A. M. James, C. Chen, W. S. Irby, E. M. Dotson, and D. K. McLain. 1993. Conspecificity of the ticks *Ixodes scapularis* and *i*. *Dammini* (Acari: Ixodidae). J. Med. Entomol. 30: 54–63.

Oliver Jr, J. 1977. Cytogenetics of mites and ticks. Annu. Rev. Entomol. 22: 407–429.

Prjibelski, A., D. Antipov, D. Meleshko, A. Lapidus, and A. Korobeynikov. 2020. Using spades de novo assembler. Current protocols in bioinformatics 70: e102.

Ronai, I., R. de Paula Baptista, N. S. Paulat, J. C. Frederick, T. Azagi, J. W. Bakker, K. C. Dillon, H. Sprong, D. A. Ray, and T. C. Glenn. 2024. The repetitive genome of the *Ixodes ricinus* tick reveals transposable elements have driven genome evolution in ticks. bioRxiv: 2024.2003.2013.584159.

Sharma, A., M. N. Pham, J. B. Reyes, R. Chana, W. C. Yim, C. C. Heu, D. Kim, D. Chaverra-Rodriguez, J. L. Rasgon, R. A. Harrell, II, A. B. Nuss, and M. Gulia-Nuss. 2022. Cas9-mediated gene editing in the black-legged tick, Ixodes scapularis, by embryo injection and remot control. iScience 25: 103781.

Shaw, D. E., and M. A. White. 2022. The evolution of gene regulation on sex chromosomes. Trends Genet. 38: 844–855.

Sonenshine, D. E., and R. M. Roe. 2014. External and internal anatomy of ticks, pp. 74–98. In D. E. Sonenshine and R. M. Roe (eds.), Biology of ticks, vol. 1. Oxford University Press, [Online].

Troughton, D. R., and M. L. Levin. 2007. Life cycles of seven ixodid tick species (Acari: Ixodidae) under standardized laboratory conditions. J. Med. Entomol. 44: 732–740.

Yuval, B., and A. Spielman. 1990. Sperm precedence in the deer tick *Ixodes dammini*. Physiological Entomology 15: 123–128.

Zhang, Z., S. Schwartz, L. Wagner, and W. Miller. 2000. A greedy algorithm for aligning DNA sequences. J. Comput. Biol. 7: 203–214.

