## Supplementary material for "Duplex PCR assay to determine sex and mating status of *Ixodes scapularis* (Acari: Ixodidae), vector of the Lyme disease pathogen"

### Supplementary materials

**Table S1.** Resequencing sample information and sequencing statistics.

**Table S2.** Triple-enzyme restriction-site associated sequencing sample information and sequencing statistics.

**Table S3.** Two male-specific loci identified in the triple-enzyme restriction-site associated sequencing dataset.

**Table S4.** Male and females accurately sexed using the molecular sexing method from colony samples and field samples.

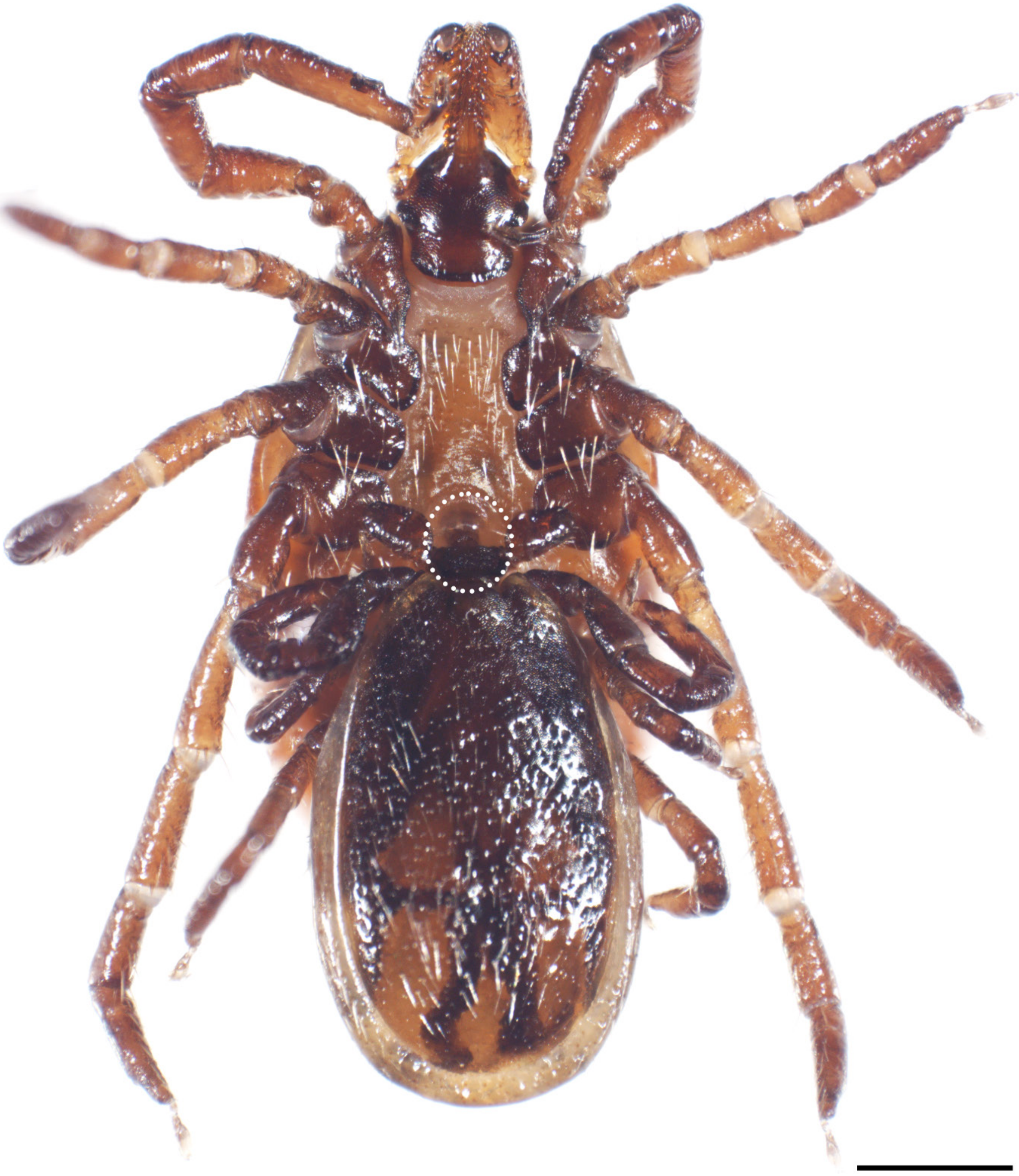

**Figure S1.** *Ixodes scapularis* mating. Ventral view of an adult male with his hypostome inserted into the adult female's genital aperture (circle) on her dorsal side. Anterior is up and scale bar represents 0.5 mm.

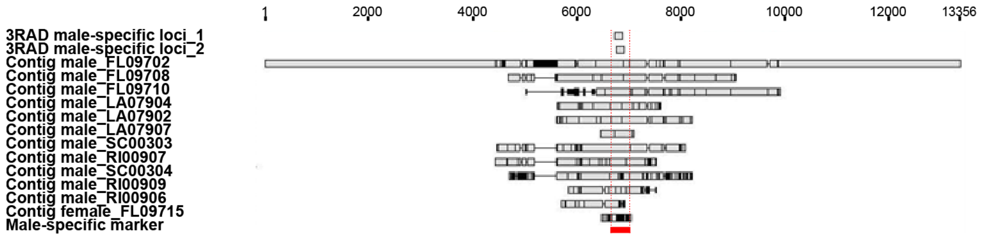


**Figure S2.** Identification of a *Ixodes scapularis* male-specific genomic region. Multiple sequence alignment: two overlapping male-specific loci from the 3RAD dataset; 11 contigs from male resequencing assemblies, with a high confidence match to the two male-specific loci; one 552 bp contig from a female resequencing assembly, with a low confidence match to the two male-specific loci; and the identified male-specific marker (red). The site position of the alignment is in bp. Nucleotide positions that match the consensus are colored grey, whereas positions that disagree with the consensus are colored black.

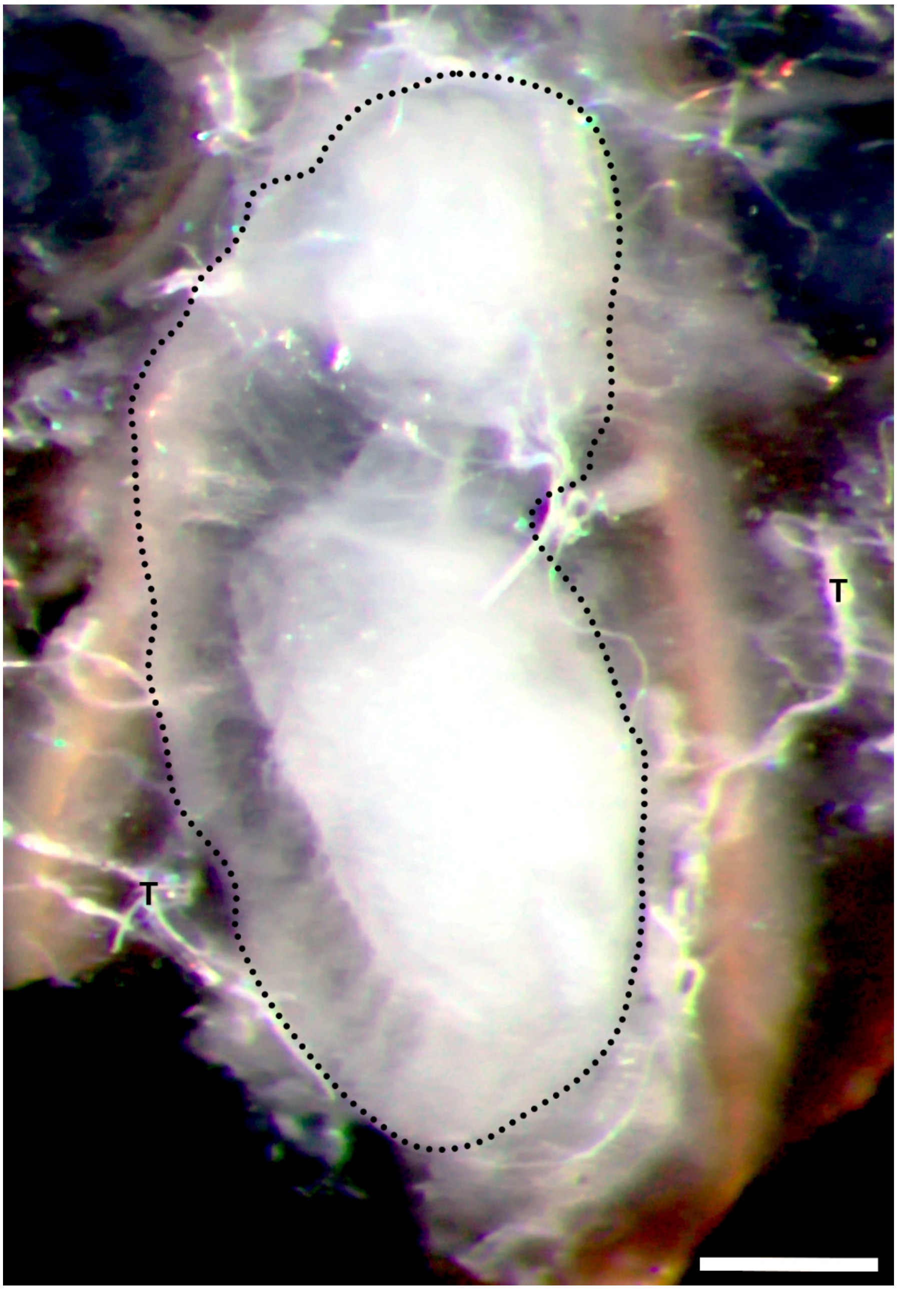


**Figure S3.** Endospermatophore (dotted line) inside a dissected adult female *Ixodes scapularis* that was paired in the laboratory mating assay. Trachea (T). Dorsal view, anterior is up and scale bar represents 0.5 mm.

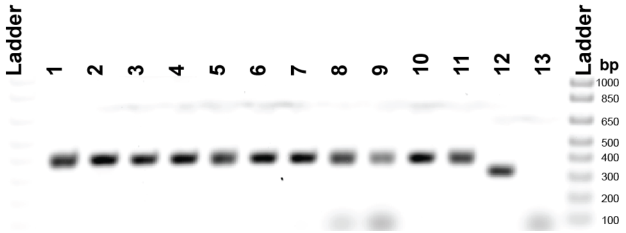


**Figure S4.** Adult female *Ixodes scapularis* that were never paired with males using the molecular sexing method (duplex PCR) visualized on an agarose gel. Primers described in Table 1. Eleven unpaired females (Lane 1-11), positive control plasmid (Lane 12) and no template negative control (Lane 13). All samples have the expected single 406 bp band and the positive control plasmid has the expected single band at 326 bp.

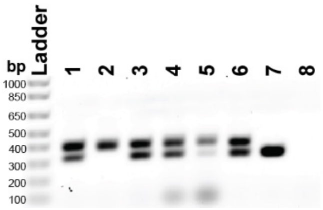


**Figure S5.** Field collected ticks that potentially mated off-host. Determination of the mating status of adult female *Ixodes scapularis* that were collected in the field using the molecular sexing method (duplex PCR) visualized on an agarose gel. Primers described in Table 1. Six females (Lane 1-6), positive control plasmid (Lane 7) and no template negative control (Lane 8). All samples have the expected single 406 bp band, however, five samples have a second band at 326 bp (Sample 1, 3, 4, 5 and 6). The positive control plasmid has the expected single band at 326 bp.
